# Single Cell Measurements and Modeling and Computation of Decision Making Errors in a Molecular Signaling System with Two Output Molecules

**DOI:** 10.1101/2023.08.22.554323

**Authors:** Ali Emadi, Tomasz Lipniacki, Andre Levchenko, Ali Abdi

## Abstract

A cell constantly receives signals and takes different fates accordingly. Given the uncertainty rendered by signal transduction noise, the cell may incorrectly perceive the signals. It may mistakenly behave as if there is a signal - although there is none, or may miss the presence of a signal that actually exists. In this paper, we consider a signaling system that has two outputs, and introduce and develop methods to model and compute key cell decision making parameters based on the two outputs, and in response to the input signal. In the considered system, the tumor necrosis factor (TNF) regulates the two transcription factors, the nuclear factor κB (NFκB) and the activating transcription factor-2 (ATF-2). These two system outputs are involved in important physiological functions such as cell death and survival, viral replication, and pathological conditions such as autoimmune diseases and different types of cancer. Using the introduced methods, we compute and show what the decision thresholds are, based on the single cell measured concentration levels of NFκB and ATF-2. We also define and compute the decision error probabilities, i.e., false alarm and miss probabilities, based on the concentration levels of the two outputs. By considering the joint response of the two outputs of a signaling system, one can learn more about complex cellular decision making processes, the corresponding decision error rates, and their possible involvement in the development of some pathological conditions.

## INTRODUCTION

Cell has to recognize and respond to the environmental variations and changes [1]. Cell fate can change based on the strength or concentration level of the extracellular stimuli or input signals. Signal transduction noise can disturb the input signals such that the cell becomes unable to correctly sense the precise concentration of different input signals and subsequently cannot properly respond. Due to the random nature of the signal transduction noise, the cellular decision is probabilistic to some extent [2]. To quantify and characterize the cell decision-making processes while incorporating their probabilistic nature, we consider a statistical signal processing approach [3]. The approach is built on decision theory and statistical signal processing concepts, to obtain optimal decision thresholds and erroneous cell decision probabilities using single cell data [3] [4]. Such a framework aims to measure the ability of the system to correctly decide on an input signal. This quantitative and probabilistic approach can be also used to characterize stochastic signaling mechanisms and phenotype induction in the context of genetic diseases [5]. One may also expand this approach to study intercellular processes together with intracellular molecular networks [6] [7].

In this paper, we consider a two-output signaling system (Fig. 1) in which the tumor necrosis factor (TNF) regulates the two transcription factors nuclear factor κB (NFκB) and the activating transcription factor-2 (ATF-2) [2]. TNF, an important antiviral cytokine, can cause noticeable damage to healthy host tissues. It is also shown that it can regulate a speed-accuracy tradeoff in the context of cell death decisions [8]. Moreover, TNF activates NFκB that leads to its nuclear translocation. NFκB is an essential gene regulator involved in cell survival, viral replication, and pathological processes such as autoimmune diseases and different types of cancer. TNF also mediates anti-apoptotic and pro-apoptotic signals and may also trigger necroptosis, as a form of pro-inflammatory cell death [9] [10]. Activation of NFκB may prevent the cell from apoptosis. The A20 (Fig. 1) mediates negative inhibitory feedback on the system input [2]. Due to this negative feedback, NFκB level may experience a reduction. With regard to the other system output, ATF-2, we note that TNF is able to activate c-Jun N-terminal kinase (JNK) pathway and stimulate the phosphorylated ATF-2 [2]. The ATF/CREB family has important physiological functions and represents a large group of basic-region leucine zipper (bZIP) transcription factors (TFs) [11]. ATFs act as heterodimers or homodimers with different bZIP transcription factors. The family includes ATF-1, ATF-2, ATF-3, ATF-4, ATF-5, ATF-6, and ATF-7 whose abilities are diversly associated with cellular processes that they regulate [11].

**Figure 1.**
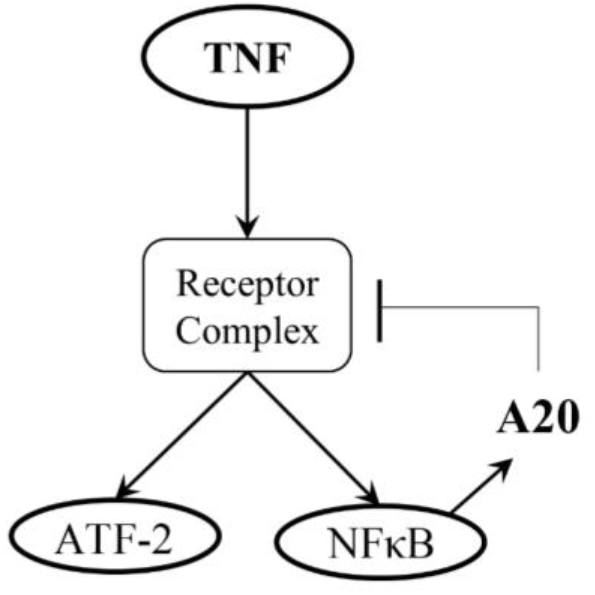
The two-output TNF— NFκB /ATF-2 signaling system.

In this paper, we aim to show how the statistical decision making framework developed for signaling systems that have only one output [3] [4], can be extended to systems with two or more outputs. More specifically and in the TNF system (Fig. 1), we compute and show what the decision thresholds are, when measurements of concentration levels of the two outputs, NFκB and ATF-2, are considered. We also define and compute decision error probabilities based on the two-output measurements and compare them with single-output decisions. By considering the joint response from the two outputs of the system, we intend to take one further step towards understanding cell decision making processes and the associated decision error probabilities.

The rest of the paper is organized as follows. First, the single cell experimental data of the TNF— NFκB /ATF-2 signaling system are introduced. Then the two-output system data and the associated decision threshold boundaries are computed and presented in graphical forms. Afterwards, various computed decision error probability results are presented and discussed, followed by detailed explanations of the computational and analysis methods. At the end, some concluding results are presented.

## RESULTS AND DISCUSSIONS

### Single Cell Data of the Two-Output TNF— NFκB/ATF-2 System

The data set was obtained from 3T3-immortalized mouse embryonic fibroblasts [2]. Nuclear concentrations of NFκB and ATF-2 were measured using immunocytochemistry of thousands of mice fibroblasts exposed to different TNF levels [2]. Further details of the used methods can be found in [12]. As explained in INTRODUCTION, decision making performance and probabilities of the two-output system in Fig. 1 - in response to its input TNF signal - are investigated in this paper, because of the high involvement of the NFκB and ATF-2 transcription factors in cell death and survival processes.

### Graphical Representation of the Two-Output System Data and the Decision Thresholds

To characterize and measure the decision probabilities on whether the TNF level is low or high, based upon the nuclear concentrations of NFκB and ATF-2, four TNF levels of 0.013, 0.082, 3.2, and 50 ng/mL are arbitrarily chosen, where the first is considered to be low TNF, and the last three are considered as high TNF concentrations. Extension to deciding on more than two input signal levels, for instance, three input signal levels, is possible, as outlined in [4]. Fig. 2 presents some important two-output graphics of cells responses after 30 minutes of TNF exposure. More specifically, panels A to C of Fig. 2 show the scatter plots of nuclear NFκB and ATF-2, where the high TNF level is 0.082, 3.2 or 50 ng/mL, respectively, compared to the fixed low TNF level of 0.013 ng/mL. The associated bivariate Gaussian probability density functions (PDFs) for nuclear NFκB and ATF-2 (see METHODS) are shown in the last row of Fig. 2. Finally, panels D to F of Fig. 2 depict the top view heatmaps of these bivariate Gaussian PDFs, along with the corresponding optimal decision threshold curves (DTCs). An optimal DTC is a curve that divides the two-dimensional parameter space, here the NFκB/ATF-2 plane, such that the decision error probability - defined later in Equation (3) -is minimized. Each optimal DTC is graphed by solving a quadratic equation obtained from the maximum likelihood decision-making principle (see METHODS). As an example, we see the optimal DTC in Fig. 2D that divides the NFκB/ATF-2 plane such that the region on its left corresponds to the low TNF decision and the region on its right relates to the high TNF decision. Details of how an optimal DTC is computed are provided in METHODS. Given the overlap between the two heatmap clusters, the decision error probability, based on the optimal DTC, is 0.245. As one can expect, as the high TNF level increases to 3.2 and 50 ng/mL (see Fig. 2E and Fig. 2F), the two heatmap clusters become separated and the decision error probability decreases to 0.05 and 0.03, respectively. These and other decision error probabilities for various scenarios are later computed and examined in the Decision Error Probabilities of the Two-Output System section.

**Figure 2.**
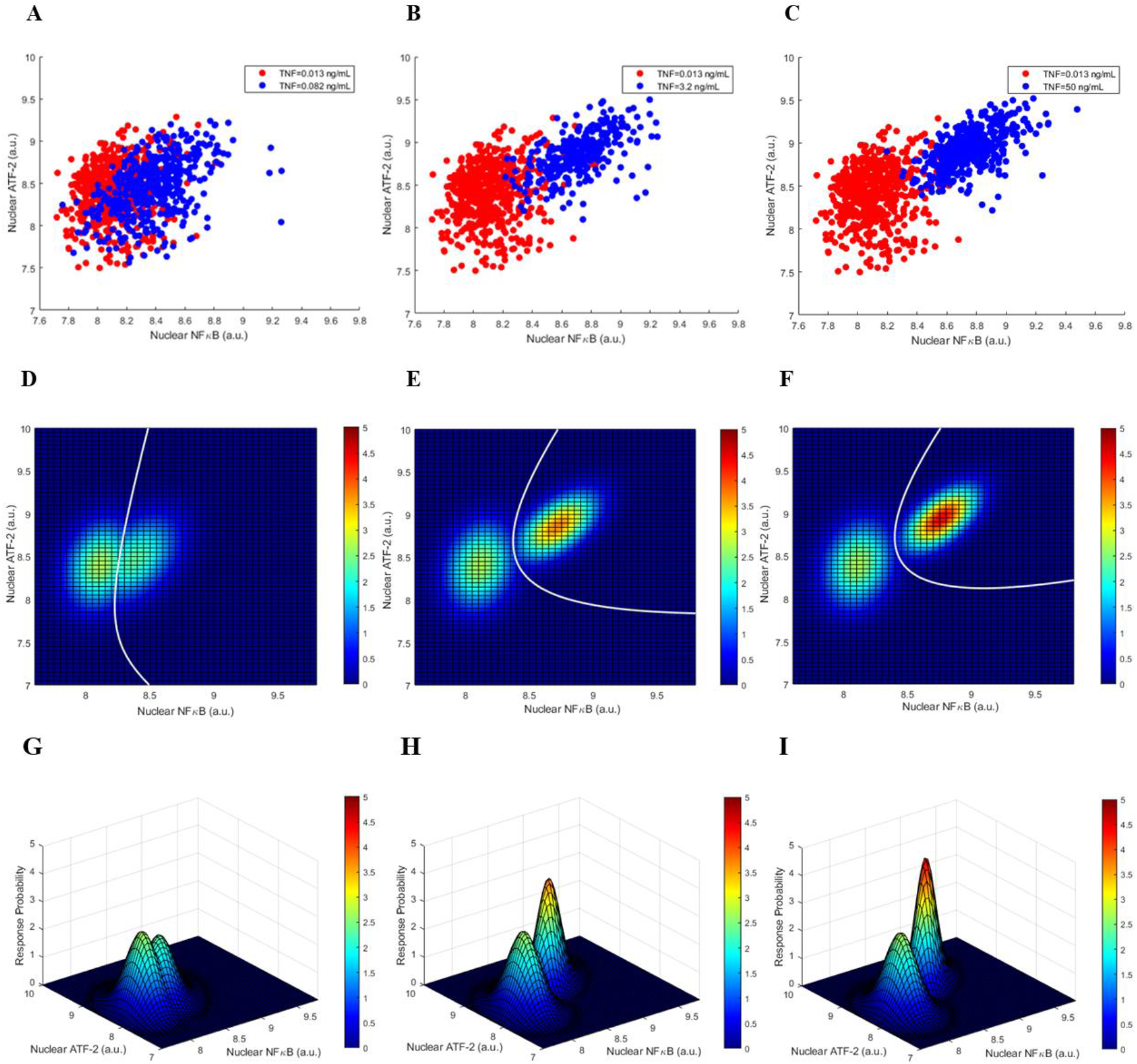
Cells responses after 30 minutes of TNF exposure. (A,B,C) Scatter plots of nuclear NFκB and ATF-2 when high TNF level is 0.082, 3.2, or 50 ng/mL, respectively. (D,E,F) Top view heatmaps of bivariate Gaussian probability density functions (PDFs) for NFκB and ATF-2, when high TNF= 0.082, 3.2, or 50 ng/mL, respectively, together with the corresponding optimal decision threshold curves (DTCs) in white. (G,H,I) Bivariate Gaussian PDFs for NFκB and ATF-2 when high TNF= 0.082, 3.2, or 50 ng/mL, respectively (The D, E and F panels are top view heatmaps of these bivariate Gaussian PDFs). In all cases, low TNF=0.013 ng/mL.

The two-output graphics of cells responses, after 4 hours of exposure to different levels of TNF, are shown in Fig. 3. We observe more overlap between the two heatmap clusters, compared to Fig. 2. This can be perhaps explained by noting that the inhibitory feedback of A20 becomes active over time, that results in reductions in nuclear NFκB and ATF-2 concentrations. The associated changes in the decision error probabilities are computed and examined in the Decision Error Probabilities of the Two-Output System section.

**Figure 3.**
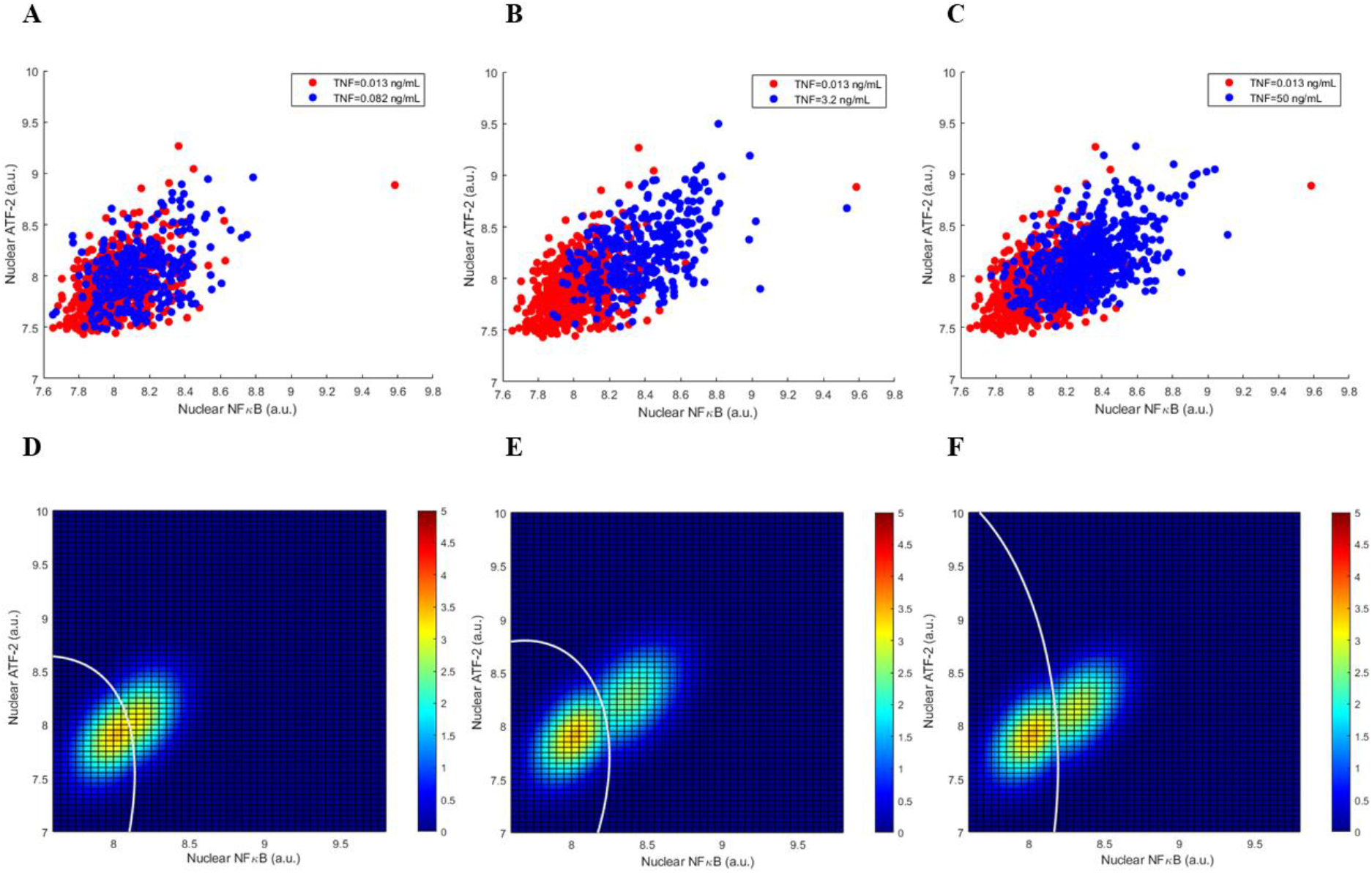
Cells responses after 4 hours of TNF exposure. (A,B,C) Scatter plots of nuclear NFκB and ATF-2 when high TNF level is 0.082, 3.2, or 50 ng/mL, respectively. (D,E,F) Top view heatmaps of bivariate Gaussian PDFs for NFκB and ATF-2, when high TNF= 0.082, 3.2, or 50 ng/mL, respectively, together with the corresponding optimal DTCs.

### Decision Error Probabilities of the Two-Output System

Information theoretical studies of single cell data of a TNF signaling system have demonstrated that a cell can distinguish between low and high TNF concentrations at the system input [2]. Due to the uncertainty caused by the signal transduction noise, two types of incorrect decisions can be made: Deciding that TNF is high while it is low indeed, or deciding that TNF is low whereas it is actually high. These two are called false alarm and miss errors, respectively [3]. In what follows, we present (Fig. 4) and discuss the computed false alarm and miss error probabilities, *P*_FA_ and *P*_M_, respectively, using measured nuclear NFκB and ATF-2 concentrations as the two outputs of the signaling system.

**Figure 4.**
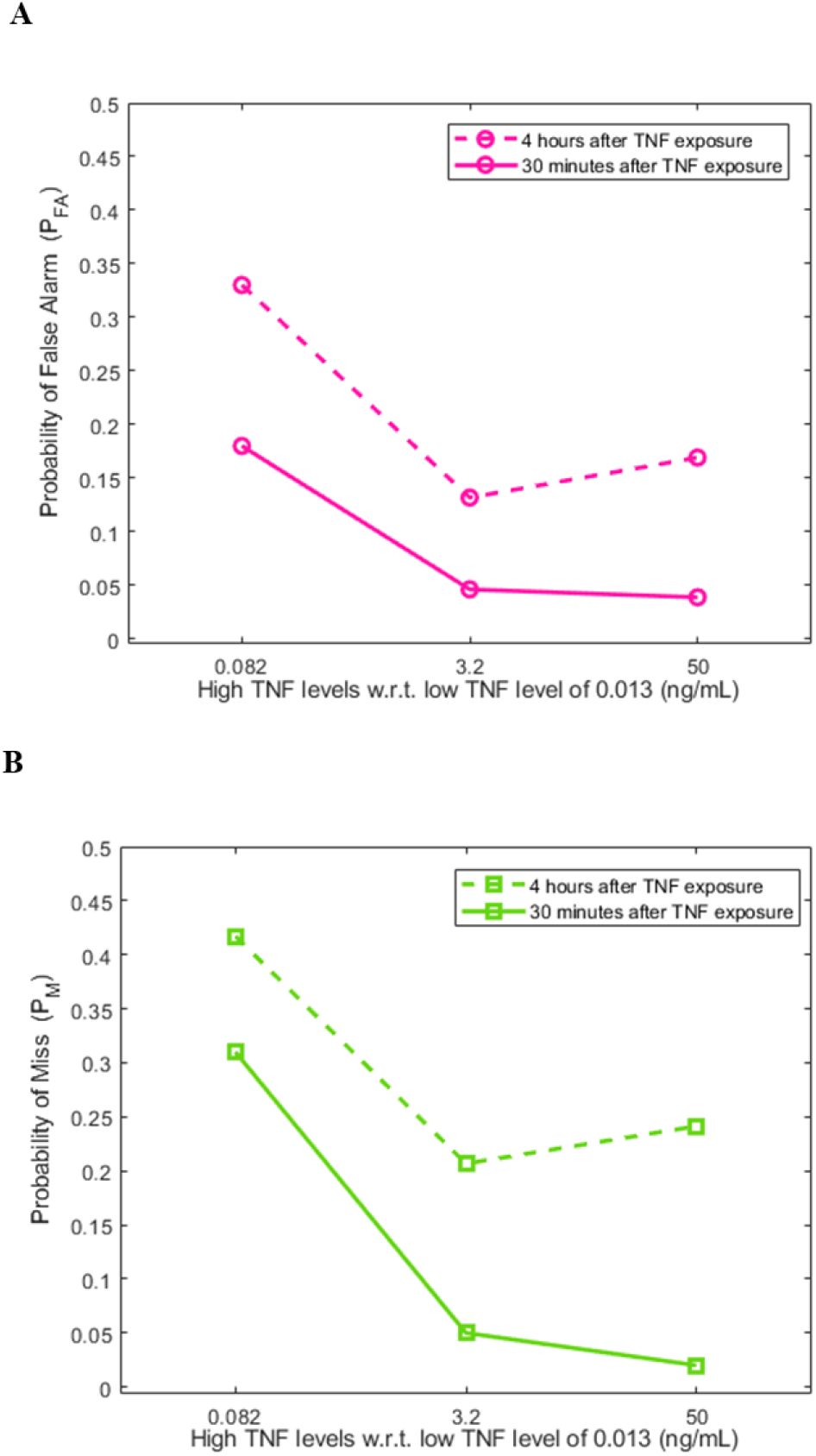
Decision error probabilities in cells based on measured nuclear NFκB and ATF-2 concentrations as the two outputs of the signaling system, after 30 minutes and 4 hours of TNF exposure. (A) Probabilities of false alarm. (B) Probabilities of miss.

To graphically explain how the *P*_FA_ and *P*_M_ error probabilities are computed, consider the high TNF level of 0.082 ng/mL in Fig. 4 as an example. The computed 0.18 false alarm probability (Fig. 4A, 30 minutes) is obtained by computing the volume under the low TNF bivariate PDF in Fig. 2G that falls on the right-hand side region of the DTC graphed in Fig. 2D - a curve that divides the NFκB/ATF-2 plane to two separate decision regions. This is the false alarm region (see METHODS). Moreover, the computed 0.31 miss probability (Fig. 4B, 30 minutes) is obtained by computing the volume under the high TNF bivariate PDF in Fig. 2G that falls on the left-hand side region of the DTC graphed in Fig. 2D. This is the miss region (see METHODS). Other false alarm and miss error probabilities in Fig. 4 are similarly computed.

We note a monotone decrease of both decision error probabilities in 30 minutes (Fig. 4), as the TNF signal becomes stronger. This can be attributed to the linear structure of the pathway in short term, when the feedback is not active yet. In 4 hours, however, we do not observe a monotone decrease of the decision error probabilities as the TNF signal strength increases (Fig. 4). This is perhaps because of the activation of the A20 feedback that renders a nonlinear structure for the pathway.

Another noteworthy observation is that the *P*_FA_ and *P*_M_ error probabilities are higher in 4 hours (Fig. 4). The two heatmap clusters exhibit more overlaps in 4 hours (Fig. 3), compared to 30 minutes (Fig. 2). This can be associated with the reductions in nuclear NFκB and ATF-2 concentrations in 4 hours, caused by the negative A20 feedback.

To investigate how much the importance of each individual system output is, when it comes to decision making using both outputs, we use the minimum redundancy maximum relevance (MRMR) algorithm [13] implemented in MATLAB®, originally developed for feature selection in classification problems. The MRMR algorithm finds an optimal set of features, so that the redundancy in the feature set is minimized, while its relevance to the response variable is maximized. The algorithm defines relevance as the mutual information between each feature and the response variable, and measures redundancy as the mutual information among the features. By defining the mutual information quotient (MIQ) parameter as the ratio of the relevance over the redundancy of a feature, the MRMR algorithm ranks the features. It also computes an importance score for each feature using a recursive approach [14]. In our case of NFκB and ATF-2, the two outputs of the system, after finding the output with the highest rank, the algorithm assigns the corresponding relevance value as the importance score of the highest-ranked output. The importance score of the second-ranked output is the importance score of the first-ranked output, multiplied by the ratio of the first-ranked output’s MIQ over the second-ranked output’s MIQ. Further details are provided in [14].

Now we compute the importance scores for NFκB and ATF-2 (Table 1), to quantify their importance in terms of their ability to render a decision on the status of the input TNF signal. Given the lower scores of ATF-2, one may say that ATF-2 possibly plays a smaller role in the decision making. This is confirmed by computing and comparing univariate decision error probabilities with the bivariate probabilities. For example, for the high TNF level of 0.082 ng/mL and 30 minute data we have these univariate miss probabilities of *P*_M_(NFκB) = 0.32 and *P*_M_(ATF-2) = 0.44, and the bivariate miss probability of *P*_M_(NFκB & ATF-2) = 0.31. We note that the bivariate probability is closer to the univariate probability rendered by NFκB only. For the same high TNF level and 4 hours data, we observe the same pattern - the univariate miss probabilities are *P*_M_(NFκB) = 0.41 and *P*_M_(ATF-2) = 0.47 - while the bivariate miss probability is *P*_M_(NFκB & ATF-2) = 0.42. A similar behavior is observed for the false alarm probability.

**Table 1.** Importance scores of NFκB and ATF-2 for decision making in cells after 30 minutes and 4 hours of TNF exposure.

| Time | High TNF level (ng/mL) | Importance score |  |
| --- | --- | --- | --- |
|  |  | NFκB | ATF-2 |
| 30 minutes | 0.082 | 0.18 | 0 |
|  | 3.2 | 0.49 | 0.15 |
|  | 50 | 0.57 | 0.2 |
| 4 hours | 0.082 | 0.05 | 0.01 |
|  | 3.2 | 0.3 | 0.1 |
|  | 50 | 0.24 | 0.07 |

## METHODS

In this paper, cellular decision making in the TNF— NFκB /ATF-2 signaling system is considered as the following hypothesis testing problem:

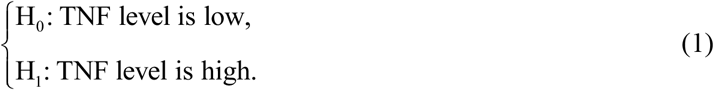

The cell may make each of these two mistakes due to the signal transduction noise: Deciding that TNF is high at the input of the system while it is actually low - declaring H_1_ while H_0_ is true - or deciding that TNF is low although in fact it is high - declaring H_0_ when H_1_ is true. These are false alarm and miss incorrect decisions, respectively, with the following probabilities:

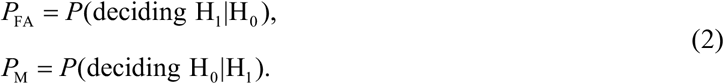

Additionally, the overall error probability *P*_E_ of making decisions is a weighted summation of *P*_FA_ and *P*_M_:

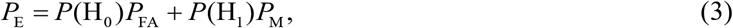

where *P*(H_0_) and *P*(H_1_) represent the prior probabilities of the hypotheses H_0_ and H_1_, respectively.

The optimal decision-making approach minimizes *P*_E_ [15]. To understand how such decisions are made, suppose **z** is the *N*-element vector of observed variables or data (in our case, we have *N* = 2 and the components of **z** represent NFκB and ATF-2). Let *p*(**z** | H_0_) and *p*(**z** | H_1_) be the conditional probability density functions (PDFs) of **z** under the hypotheses H_0_ and H_1_, respectively. The optimal decision rule is derived from the maximum likelihood principle [15] that chooses the hypothesis with the highest likelihood of occurrence as the best - optimal - decision. More specifically, the optimal decision rule compares the conditional likelihood ratio *L*(**z**) = *p*(**z** | H_1_) / *p*(**z** | H_0_) with the likelihood threshold of *γ* = *P*(H) / *P*(H) and decides H_1_ if *L*(**z**) > *γ*, and decides H_0_ otherwise. In general, *L*(**z**) = *γ* represents the optimal decision threshold hypersurface and for *N* = 2, it represents the optimal decision threshold curve (DTC).

To evaluate the performance of this optimal decision rule, false alarm and miss probabilities [15] need to be computed using the following *N*-variate integrals:

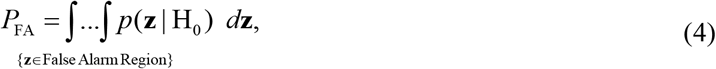

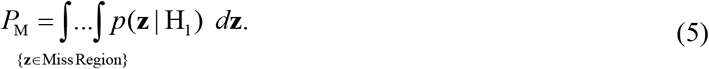

As discussed later, Gaussian PDFs can represent the data. An *N*-variate Gaussian PDF for **z** under the *i*-th hypothesis H_*i*_ can be written as [15]:

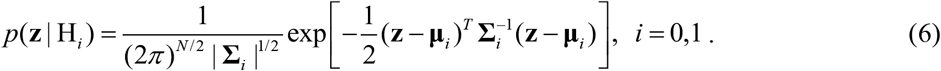

In the above equation, **μ**_*i*_ and **Σ** _*i*_ are the mean vector and the covariance matrix of **z** under H_*i*_, respectively, | **Σ**_*i*_ | and 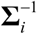 represent the determinant and the inverse of the matrix **Σ** _*i*_, respectively, and ^*T*^ denotes the transpose operation.

### A. Using the Likelihood Ratio to Compute the Optimal Decision Thresholds and the Decision Error Probabilities for the TNF— NFκB/ATF-2 System

To compute the optimal decision thresholds and the decision error probabilities using the likelihood ratio, first we present the univariate methods that have simpler equations, followed by the bivariate methods.

### A1. Univariate Decision Analysis

We use the nuclear NFκB and ATF-2 concentrations of thousands of cells that are exposed to the TNF concentrations of 0.013, 0.082, 3.2, and 50 ng/mL, after 30 mins and 4 hours [2]. For the univariate decision analysis, let us define *x* = ln(Nuclear NFκB), where ln is the natural logarithm. Examination of the data reveals that a Gaussian PDF can be used to model the *x* variable:

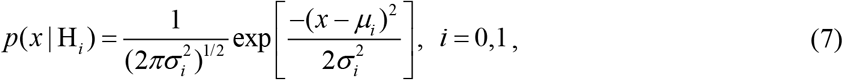

where *μ*_*i*_ and 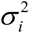 are the mean and the variance under the *i*-th hypothesis H_*i*_, respectively. The TNF level under H_0_ is 0.013 ng/mL, while under H_1_ it is 0.082, 3.2 or 50 ng/mL.

The optimal likelihood-based decision rule decides H_1_ if:

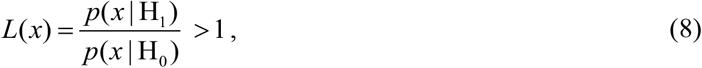

for equi-probable hypotheses. To obtain the optimal decision threshold for *x*, the following *p*(*x* | H_1_) = *p*(*x* | H_0_) equation needs to be solved for *x*:

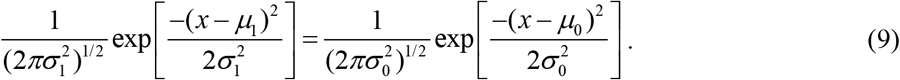

After algebraic simplification of Equation (9), the following quadratic equation is obtained:

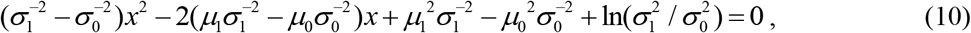

and by solving this equation numerically, the optimal decision threshold can be computed.

The false alarm and miss probabilities can be computed using the following formulas [3]:

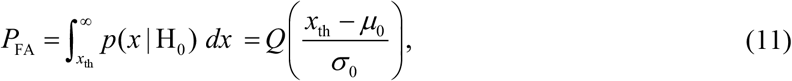

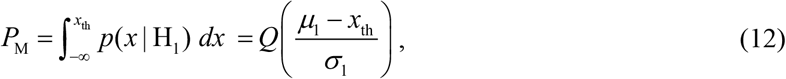

where the *Q* function is defined below:

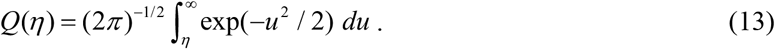

By defining *y* = ln(Nuclear ATF-2), all the above equations and results can be used, after replacing *x* in there with *y*.

### A2. Bivariate Decision Analysis

For the bivariate decision analysis, we consider *x* = ln(Nuclear NFκB) and *y* = ln(Nuclear ATF-2), and define the two-element vector **z** =[*x y*]^*T*^. Upon substituting *N* = 2 in Equation (6), the following bivariate PDF can be written for **z**:

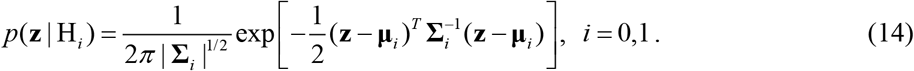

Here **μ** _*i*_ and **Σ**_*i*_ are the mean vector and the covariance matrix under the *i*-th hypothesis H_*i*_, respectively:

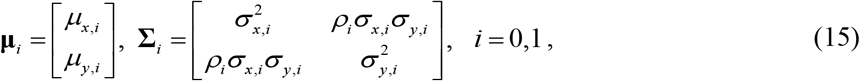

where *ρ*_*i*_ is the correlation coefficient between *x* and *y* under the *i*-th hypothesis H_*i*_.

The optimal likelihood-based decision rule decides H_1_ if:

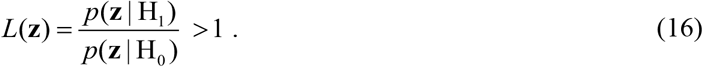

To find the optimal DTC in the *x*-*y* plane, the following in *p*(**z** |H_1_) = *p*(**z** |H_0_) equation needs to be solved terms of *x* and *y*:

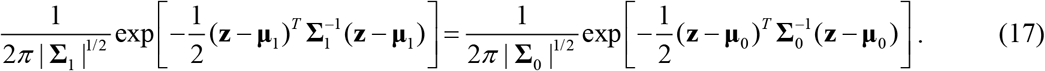

By taking the natural logarithm of both sides of Equation (17) and rearranging some terms, the following bivariate quadratic equation is obtained:

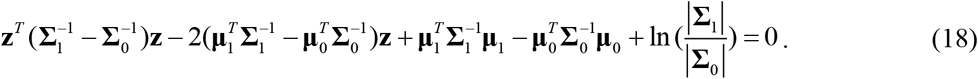

By solving this equation numerically, the optimal DTC can be computed and graphed in the *x*-*y* plane. For

*N* = 1 and when **z** includes only the one variable *x*, Equation (18) reduces to (10).

The false alarm and miss probabilities can be computed using Equations (4) and (5) with *N* = 2, respectively:

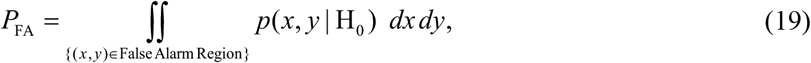

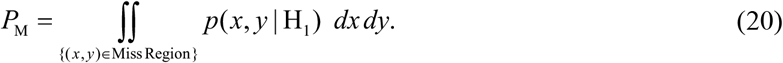

The bivariate integrals in Equations (19) and (20) are computed using Monte Carlo integration.

### B. Using the Discriminant Function to Compute the Decision Error Probabilities for the TNF— NFκB/ATF-2 System

Here we explain how to use the discriminant function [4] [16] to compute the decision error probabilities, without computing multivariate integrals. The discriminant function for the *i*-th hypothesis H_*i*_ is defined as:

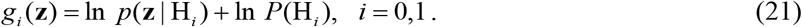

By substituting Equation (6) together with *N* = 2 in (21) we obtain:

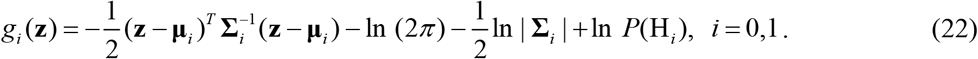

The optimal decision rule decides H_1_ if *g*_1_(**z**) > *g*_0_ (**z**).

To compute the false alarm probability *P*_FA_ using the discriminant functions and the **z** data points available under H_0_, we compare numerical values of *g*_1_ (**z**) and *g*_0_ (**z**) for each **z** under H_0_, count the number of times that *g*_1_(**z**) > *g*_0_ (**z**), that is, when the false alarm event occurs, and then divide it by the number of the **z** data points available under H_0_. The miss probability *P*_M_ is similarly computed.

## CONCLUSION

In this paper, we introduce and develop a set of statistical signal processing and decision theoretic methods and metrics for modeling and measurement of decision-making errors in a TNF signaling system that regulates the two important transcription factors NFκB and ATF-2. As a useful and informative visual tool, first optimal decision threshold curves (DTCs) are computed and graphed in the two-dimensional NFκB/ATF-2 plane (Fig. 2 D,E,F, and Fig. 3 D,E,F), where an optimal DTC is a curve that divides the two-dimensional output space such that the error probability of making decisions on the TNF signal level is minimized.

Second, using measured nuclear NFκB and ATF-2 concentrations, false alarm (*P*_FA_) and miss (*P*_M_) error probabilities are computed. Here *P*_FA_ is the error probability of deciding that TNF is high while it is indeed low, whereas *P*_M_ is the error probability of deciding that TNF is low, even though it is actually high. We observe a monotone decrease of both decision error rates in 30 minutes versus the TNF signal level (Fig. 4), perhaps because of the linear structure of the pathway in short term. In 4 hours, however, a monotone decrease of the decision error probabilities is not observed (Fig. 4), possibly due to the activation of the A20 feedback that induces a nonlinear structure for the pathway. We also notice that *P*_FA_ and *P*_M_ increase from 30 minutes to 4 hours (Fig. 4A and Fig. 4B, respectively). This can be because of the reductions in the nuclear NFκB and ATF-2 concentrations in 4 hours - due to the negative A20 feedback - that make the two heatmap clusters get closer to each other and overlap further (Fig. 3), compared to the 30 minute heatmap clusters overlap (Fig. 2).

Third, we look at each system output alone, to understand their relative individual importance in providing decisions on the status of the input TNF signal, compared to the decisions made using both outputs together. We observe that ATF-2 plays a smaller role, compared to NFκB. This behavior of one output being less important than the other, however, may be specific to the signaling system studied here, and may not necessarily hold true for other signaling systems.

In conclusion, the developed statistical signal processing and decision-theoretic metrics and methods can quantify complex cellular decision making processes and behaviors. The introduced metrics and methods can be applied to other and larger signaling systems that have several inputs such as ligands or second messengers and outputs such as multiple transcription factors.

## ACKNOWLEDGMENTS

Tomasz Lipniacki was supported by the National Science Centre (Poland) [grant number 2019/35/B/NZ2/03898], and Andre Levchenko was supported by NSF [grant number 2231765 from the MCB division].

